# Antiretroviral Adverse Drug Reactions Pharmacovigilance in Harare City, Zimbabwe, 2017

**DOI:** 10.1101/358069

**Authors:** Hamufare Mugauri, Owen Mugurungi, Tsitsi Juru, Notion Gombe, Gerald Shambira, Mufuta Tshimanga

## Abstract

**Introduction:** Key to pharmacovigilance is spontaneously reporting all Adverse Drug Reactions (ADR) during post-market surveillance. This facilitates identification and evaluation of previously unreported ADR’s, acknowledging the trade-off between benefits and potential harm of medications. Only 41% ADR’s documented in Harare city clinical records for January to December 2016 were reported to Medicines Control Authority of Zimbabwe (MCAZ). We investigated reasons contributing to underreporting of ADR’s in Harare city.

**Methods:** A descriptive cross-sectional study and the updated Centers for Disease Control (CDC) guided surveillance evaluation was conducted. Two hospitals were purposively included. Seventeen health facilities and 52 health workers were randomly selected. Interviewer-administered questionnaires, key informant interviews and WHO pharmacovigilance checklists were used to collect data. Likert scales were applied to draw inferences and Epi info 7 used to generate frequencies and proportions.

**Results:** Of the 52 participants, 32 (61.5%) distinguished the ADR defining criteria. Twenty-nine (55.8%) knew system’s purpose whilst 28 (53.8%) knew the reporting process. Knowledge scored average on the 5-point-Likert scale. Thirty-eight (73.1%) participants identified ADR’s following client complaints and nine (1.3%) enquired clients’ medication response. Forty-six (88.5%) cited non-feedback from MCAZ for underreporting. Inadequate ADR identification skills were cited by 21 (40.4%) participants. Reporting forms were available in five (26.3%) facilities and reports were generated from hospitals only. Forty-two (90.6%) clinicians made therapeutic decisions from ADR’s. Averaged usefulness score was 4, on the 5-point-Likert scale. All 642 generated signals were committed to Vigiflow by MCAZ, reflecting a case detection rate of 4/ 100 000. Data quality was 0.75–1.0 (WHO) and all reports were causally assessed.

**Conclusion:** The pharmacovigilance system was useful, simple, and acceptable despite being unstable, not representative and not sensitive. It was threatened by suboptimal health worker knowledge, weak detection strategies and referral policy preventing ADR identification by person place and time. Revisiting local policy, advocacy, communication and health worker orientation might improve pharmacovigilance performance in Harare city.

## Introduction

Pharmacovigilance (PV) is the practice of monitoring the effects of medical drugs after they have been licensed for use, in order to identify and evaluate previously unreported adverse drug events (ADE) and reactions (ADR) (1). This is in recognition of the trade-off between the benefits and the potential harm of all medications (2). Rapidly increasing antiretroviral therapy (ART) access globally, has transformed HIV infection into a chronic, manageable condition with prolonged survival times (3). Consistent with typical chronic therapy, drug-related toxicities remain a major challenge in resource-constrained settings due to a limited formulary for mitigation and inadequately trained personnel (4). Treatment-limiting drug toxicities are resulting in an added layer of complexity in the management of HIV by impairing patient adherence to treatment, leading to inferior clinical outcomes and higher cost to the public health system (5).

The Medicines Control Authority of Zimbabwe (MCAZ) which houses the National Pharmacovigilance Centre, derives its mandate from the Medicines and Allied Substances Control Act (MASCA), Chapter 15:03, enacted in 1997 (6). This legislation provides the impetus for MCAZ’s stewardship role in regard to medicines licensure and regulation in the country. The main thrust being ensuring improved patient care and safety during medical and paramedical interventions, thereby improving public health and safety in relation to the use of medicines. In addition, the system promotes understanding, education and clinical training in pharmacovigilance and its effective communication to the public (7). The operations of the Centre are guided by WHO guidelines for setting up and running a national pharmacovigilance Centre. In this regard, the Zimbabwe National Pharmacovigilance Policy and Guidelines serve as a handbook for pharmacovigilance activities in the country (8).

The bedrock of pharmacovigilance systems, that aim to improve medicinal products safety, is prompt, spontaneous reporting of adverse drug reactions (ADRs) as a key step to their mitigation as well as updating the drug information database (9–11). It is, therefore, a mandatory requirement for health care providers to timely report all suspected and confirmed ADRs. This is particularly imperative in Zimbabwe, where the treat all strategy is being implemented, since June 2016, and has resulted in the number of people on HIV treatment rapidly increasing (12).

A preliminary review of ADR data for ARV’s from Harare City which was reported through the MCAZ and through Opportunistic Infections (OI) records, captured between 01 January and 31 December 2016 was conducted. A 41% discrepancy was discovered in these two reporting systems, with more cases appearing in OI records than what was reported to MCAZ (6). This indicated poor reporting practices that impede accurate quantification of the prevalence of ADR’S. Failure to detect and report adverse drug reactions compromises patient safety and results in missed opportunities to update drug safety profiles. It is within this background that we evaluated the ADR surveillance system in Harare City in order to identify the reasons for underreporting and recommend solutions.

**Figure 1:**
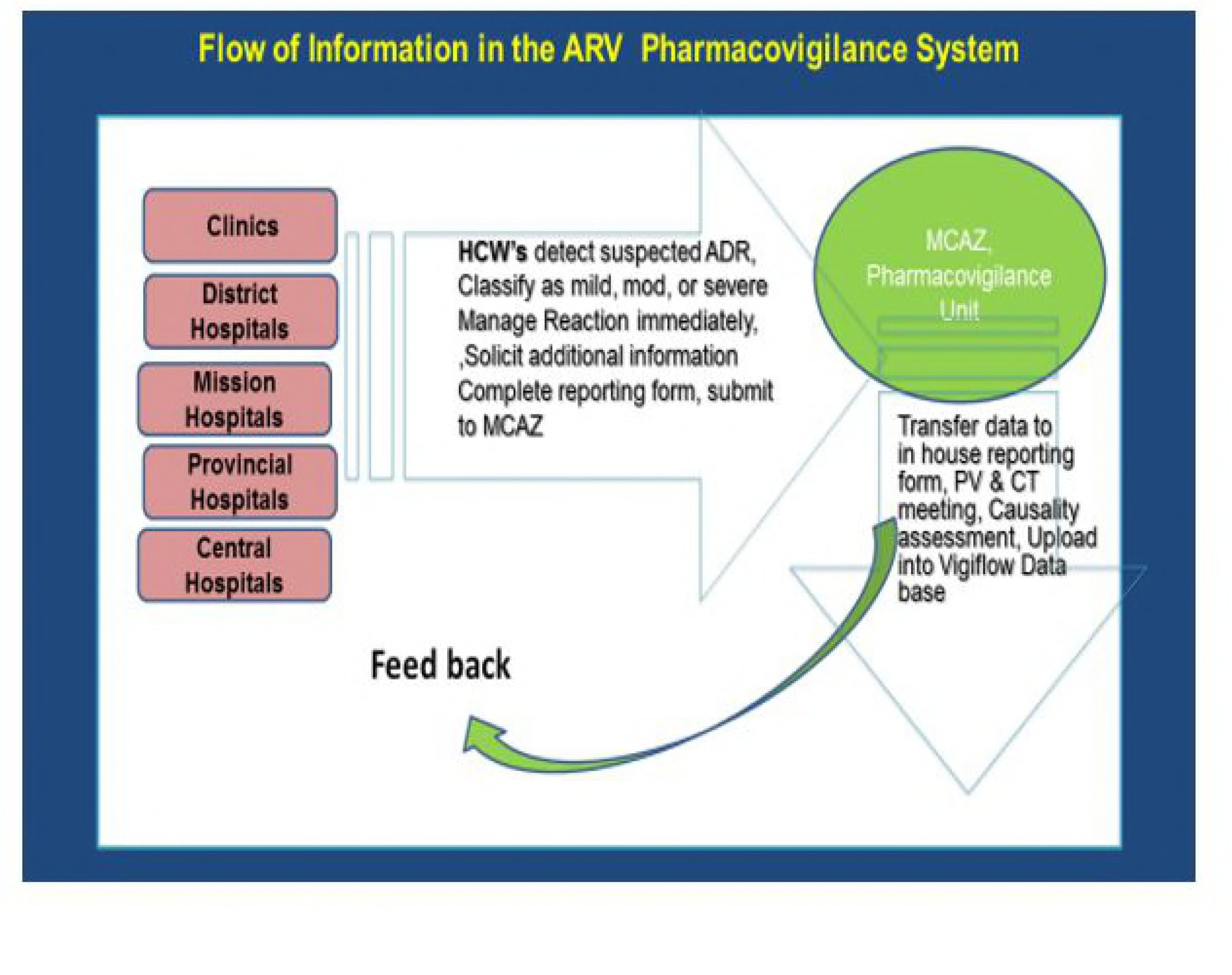
The ARV ADR Surveillance Flow Diagram

When an ADR case is suspected or confirmed, an in-house reference number is assigned. The data collected and entered into the standard reporting form should be checked for completeness. Additional information and clarifications should be solicited from the reporter before the report is filed. Once done, a completed form should be submitted to MCAZ within 14 days for spontaneous reporting (SR).

Meanwhile, corrective interventions will be of course. At MCAZ, received reports are transferred to the MCAZ reporting form to be tabulated for causality assessment at the next seating of the pharmacovigilance and clinical trials (PVCT) meeting. The causality assessment process involves analysis of the reaction against a set of key aspects that include the strength of the association, consistency of the observed evidence, temporality, dose-response and identification of possible confounders [13].The recommendations derived from this meeting are then implemented, which may be a request for further information where clarity is desired and informing healthcare facilities of findings. The data is also uploaded into Vigiflow database, including causality assessment outcome and case summary reports.

## Materials and Methods

We conducted a descriptive cross-sectional study and surveillance system evaluation using updated CDC guidelines for surveillance system evaluation as a mixed method. Health Personnel involved in the ARV-ADR surveillance system were randomly selected to participate in the evaluation. These included doctors, pharmacists, nurses and pharmacy technicians. Harare City’s two hospitals were purposively selected for the study and seventeen out of 38 clinics were randomly selected for the study. At the hospitals, all available health workers (nurses, pharmacists and doctors) working in OI clinics were recruited as study participants. The sister in charge, pharmacist and doctor at the OI clinics and hospitals were purposively recruited for the study. From the clinics, nurses who were found on duty on the day of data collection were selected for the study. All data records on reported ART ADR’s for the period under study were reviewed at MCAZ and triangulated with data from the facilities.

Using Dobson formula: 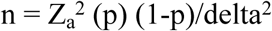, where Z_a_=1.96, p=0.5, assuming that 50% of health workers interviewed had adequate knowledge, at 20% precision and 80% power, a sample size of 47, adjusted for 10% non-response rate, sample size of 52 was reached.

A pre-tested interviewer-administered questionnaire was used to interview the health workers to determine their knowledge of the operations and usefulness of the surveillance system. The variables assessed on health worker knowledge included: the ability to accurately enumerate the key elements of an ADR, (a noxious response, unintended, at therapeutic dosage), sequentially relating the entire ADR reporting process, the purpose and the role of MCAZ in ADR Surveillance. The quality of the data generated was scored in relation to completeness, consistent with WHO evaluation criteria.

A checklist was used to assess the system’s stability. Records of all patients who were attended at the health facilities were reviewed to check on the number of ARV ADR cases documented and the number captured by the surveillance system and how many were missed. All notification forms from January to December 2016 were reviewed. Simplicity, data quality, completeness, acceptability, sensitivity, timeliness and representativeness of the system were evaluated. Epi InfoTM was used to compute frequencies, means, and proportions. The checklist for PV indicators was evaluated according to WHO score values.

Permission to carry out the study was obtained from the Institutional Ethical clearance boards for the Medicines Control Authority of Zimbabwe (MCAZ), Harare city and Ministry of Health and Child Care, Written informed consent was obtained from key informants.

## Results

### Demographic characteristics

The study successfully recruited 52 Health workers as study participants, yielding 100% response rate. Of the 52 participants recruited, 73% (n=38) were females. The majority (75%) of the participants were Registered General Nurses (RGNs). The median years of service of all participants were 9years (Q_1_= 7, Q_3_= 12). **(Table 1.)**

**Table 1.** Demographic Characteristics of Study Participants, Harare City, Zimbabwe, 2017

| Characteristics | Frequency n (%) | n=52 |
| --- | --- | --- |
| **Gender** |  |  |
| Female | 38 (73) |  |
| Male | 14 (27) |  |
| **Designation** |  |  |
| Medical Doctors | 4(7) |  |
| Registered General Nurses (RGNs) | 39 (75) |  |
| Pharmacy Technicians | 3 (6) |  |
| Primary Counsellors (PC) | 6(12) |  |
| **Median Years in Service** | 9 (<u>Q<sub>1</sub></u>= 7 , <u>Q<sub>3</sub></u>= 12 ) |  |

RGN: Registered General Nurse
PC: Primary Counsellor
Q<sub>1</sub>: First Quartile
Q<sub>3</sub>: Third Quartile

### Health Worker Knowledge of ARV ADR Surveillance

Varied proportions of respondents gave accurate responses to each variable assessed on health worker knowledge. The total score was then rated using a 5-point Likert scale which ranged from very poor, poor, fair, and good to very good. Overall, knowledge was rated as fair.

## System Attributes

### Data Quality

Data quality obtained a score range of .75–1.0, according to WHO derived from country records that were committed to WHO Vigiflow. Observed completeness of available forms from the sites was consistent with the national score.

### Simplicity

Out of the 52 participants, only 12 (29.4%) had ever completed an ADR form. Reported average time taken to complete AR forms, by those who had done so before was 14 minutes. However whilst being timed, participants took an average of 7–9 minutes 7/12 (19.6%) participants reported that the forms were easy to complete, and 10 out of 12 accurately outlined the entire reporting process for ADR’s. Forty-three (82.7%) stated that they needed formal training to be able to fill the notification forms. All ADR cases were referred to Wilkins and Beatrice road hospitals were reports are generated and submitted.

### Acceptability

90.4% of the participants felt that it was their duty to complete the ADR forms and 92.3% participants were willing to continue participating in the ADR surveillance. Thus based on the subjective assessment gathered from the interview, on average, ADR surveillance is 91.4% acceptable to health workers in Harare City.

### Stability

Twenty-one (40.4%) of the participants reported that they had ADR case definitions in their Health facilities. However only two out of 19 (10.5%) health facilities had the ADR case definition displayed. Five (26.3%) health facilities had ADR forms available in their workstations. Thirteen (25%) of the participants knew about the 2016 invented online reporting facility, but none had ever used it due to computer, Internet and knowledge challenges. One health facility (Wilkins hospital) had accessible, facility-level ADR record. All facilities had a working phone for communication.

### Usefulness- Perceptions of ADR Surveillance System, Harare City, 2017

Overall 69.2% of the participants used ADR data in patient management whilst 13.5% said they held review meetings for ADR’s. There was no evidence of minutes to the referred meetings. Clinicians made therapeutic decisions using ADR data, such as switching to next line regimen. Applying the 5 points Likert scale on the resultant usefulness score, ADR pharmacovigilance was somewhat useful with an average score of 64.5%. **(Table 2.)**

**Table 2.** Usefulness of the ARV Pharmacovigilance System, Harare, 2017

| **Variable** | **Doctors (%)** | **Nurses (%)** | **Pharmacy Technicians (%)** | **Primary Counsellors (%)** |
| --- | --- | --- | --- | --- |
| Data used in patient management (Yes) | 4 (100) | 28 (71.8) | 1 (33.3) | 3 (50) |
| ADR Meetings held (Yes) | 0 | 5 (12.8) | 0 | 2 (33.3) |
| Decisions based on ADR's (Yes) | 4 (100) | 38 (97.4) | 3 (100) | 4 (66.7) |
| Thought ADR Surveillance is useful | 4 (100) | 39 (100) | 3 (100) | 4 (66.7) |
| **Overall Usefulness Rating** | 75% | 70.5% | 58.3% | 54.2% |

ADR: Adverse Drug Reaction

### Representativeness

The system was not representative. The City Council imposed protocol of referring ADR’s to their two hospitals results in an overestimation of reports generated by the hospitals, at the same time underestimating the prevalence of ADR’s within the community health facilities by person place and time. Many ADR’s are not being reported for fear of writing reports as required by the City health department.

### Timeliness of the ARV ADR Surveillance System in Harare City, 2017

Severe and Moderate reactions were all (100%) reported to the authority on time (within 48 hours), entirely from the two hospitals. Mild and Incidental reactions were all (100%) treated according to facility protocol before completion and submission of the forms within 14 days.

### The sensitivity of the ADR Surveillance System in Harare City, 2017

Individual Case Safety Reports (ICSRs) received by MCAZ amounted to 642, reflecting a case detection rate of 5/ 100 000, calculated using the national population of 14 million in 2015. All received reports were tabled at PV and clinical trials committee meetings and feedback submitted to the city health authorities. We observed that e86% of the Targeted Spontaneous Reports (TSR) received since 2012 and authenticated by MCAZ were committed to the WHO Vigibase, termed Vigiflow as at 31 December 2016. Notably, 10 cases of product defects were reported in 2016, seven of which were subsequently recalled by the authority. More than 1900 adverse drug reaction cases, termed signals, were reported and the most common ones included gynecomastia, drug-induced liver injury, steven johnson syndrome, lipodystrophy and renal toxicity.

We further observed that 119 health facilities, countrywide, actively reported ADR’s (Sept 2012 to Dec 2016), yet only 32 of these facilities submitted ADR reports in 2015. This indicator is qualified by submission of ≥ 10 reports annually to the pharmacovigilance centre. The pharmacovigilance unit met the minimum requirements of a regulatory authority, according to WHO standards. **(Table 3.)**

**Table 3.** ADR Surveillance Sensitivity, Harare City, 2017

| Pharmacovigilance indicator | Key Attribute Assessed | Response and Value |
| --- | --- | --- |
| CP1 | Total number of ADR reports received in 2015 | ICSRs(AEFIs, TSR, SAEs) received by the MCAZ = 642 |
| CP1 a | Total number of ADR reports received in the previous year per 100 000 people in the population | 5/100 000 |
| CP2 | Number of reports (total @ 31/12/16) in the national database? | AEFIs = 331, TSR = 1563, SAEs = 361 |
| CP3 | Percentage of total annual reports acknowledged/issued feedback? | 100% |
| CP4 | Percentage of total reports subjected to causality assessment in the past year? | 100% |
| CP5 | Percentage of above committed to the WHO database? | 86% of TSR reports received since Sept 2012 committed to VigiFlow |
| CO1 | Signals generated in the past 5 years by the pharmacovigilance center? | +1900 ADR's |
| CO2 | Regulatory actions were taken in the preceding year | 7 products were recalled |
| P1 | Percentage of health-care facilities that had a functional pharmacovigilance unit (i.e. submits $\geq 10$ reports annually to the pharmacovigilance center)? | Sept 2012 to Dec 2016 = 119 health facilities<br>2015 = 32 health facilities submitted ADR reports. |

CP: WHO Core Pharmacovigilance Indicator
CO: Core Outcome Indicator
P1: Complementary Process Indicators
ICSR: Individual Case Safety Report
AEFI: Adverse Event Following Immunization
TSR: Targeted Spontaneous Reporting System for Adverse Reactions
SAE: Serious Adverse Events

### ARV ADR detection Strategies in Place, Harare City, 2017

Thirty-eight (73.1%) participants indicated that they detected ADR’s following client complaints, whilst 21(40.4%) identified ADR’s during clients routine review visits and examinations. Nine (17.3%) enquired clients how they were responding to treatment, whereas 34(65.1%) only identified ADR’s following clients’ failure to tolerate treatment and have defaulted.

### Reasons for underreporting of ARV ADR’s, Harare City, 2017

Whereas MCAZ is mandated to feedback on outcomes of all reported ADR’s, 46(88.5%) of the participants stated that None response by MCAZ to reported ADR’s was the reason for under-reporting of ARV ADR’s. Unavailability of reporting forms was cited 44, (84.6%) whilst 33 (63.5%) thought weak incident detection strategies was the reason for under-reporting. **(Table 4.)**

**Table 4.**
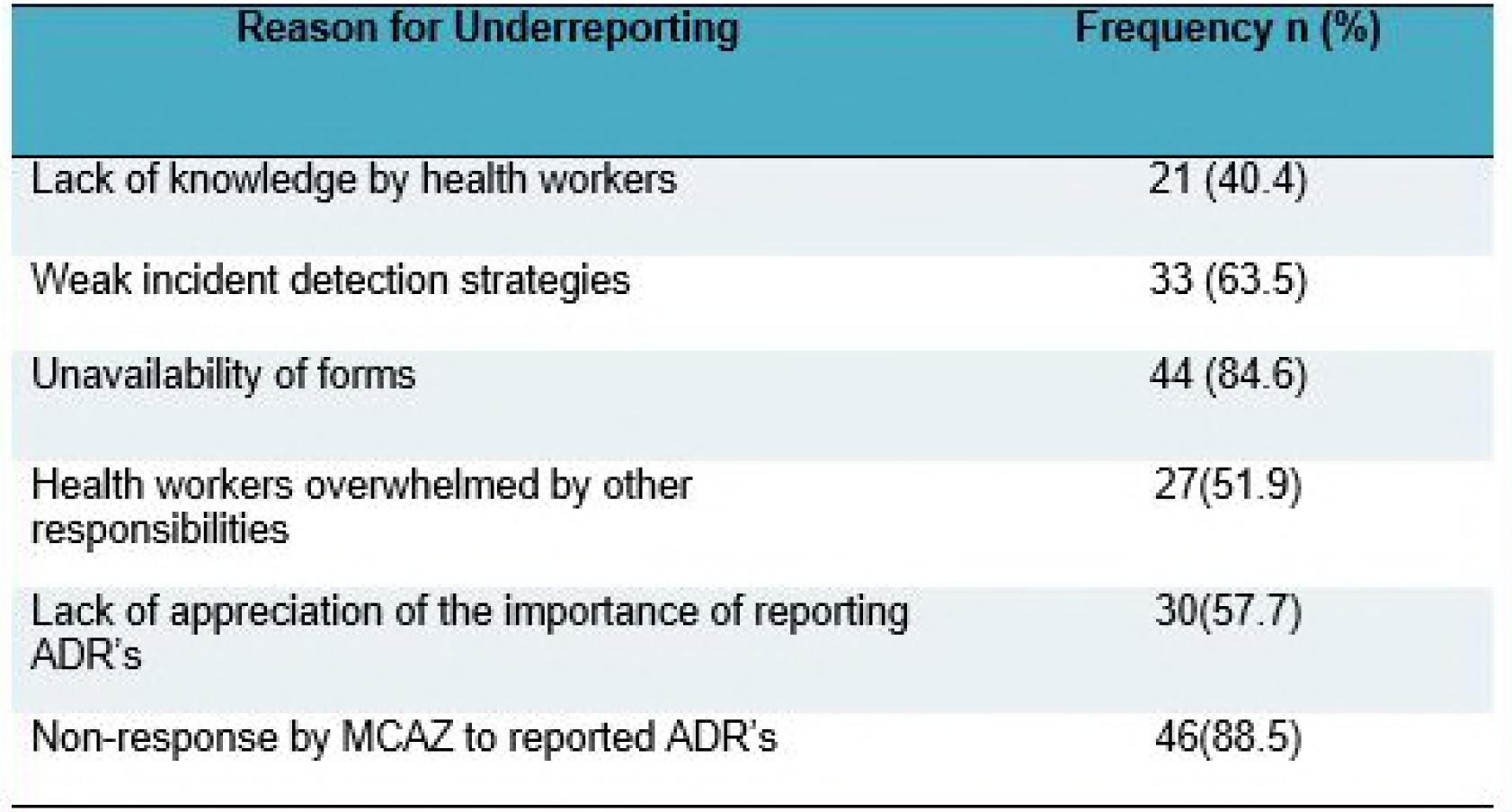

## Discussion

The data generated from the few reporting sites was of good quality in regard to completeness. A score of 0.75–1.0 for a country with a pharmacovigilance system that is still under development is remarkable. This is contrary to findings by Nderitu et al (2011) in Kenya who found incomplete records as a major hindrance to causality assessments in a developing pharmacovigilance systems (3).

Health workers are supposed to be knowledgeable about the surveillance system so that they are able to identify and investigate suspected cases during their routine conduct of duty. Knowledge on ARV pharmacovigilance surveillance in regard to qualifying ADR’s, the reporting process and the role of MCAZ was low in Harare City. This was despite their recognition that reporting ADR’s is within their scope of practice, accepting the responsibility. It was noted that the majority of the health workers were not trained (formally or on-the-job) on ARV, ADR surveillance although pharmacovigilance is a component of ART management, for which all health workers are oriented on. Poor knowledge contributed to the poor performance of the system.

The City Council imposed protocol of referring all suspected ADR’s instead of reporting directly to MCAZ resulted in it being impossible to assess the incidence and prevalence of ARV ADR’s by person place and time. All reports were being generated from Wilkins and Beatrice road hospitals.

The ARV ADR’s surveillance system in Harare city is simple. The few who filled the forms encountered no challenges. When MCAZ received suspected ADR reports, a pharmacovigilance and clinical trials committee sat to discuss and recommend causality assessment, particularly for peculiar reactions. All reports were responded to through the city health directorate for communication to reporting sites. However, health workers from reporting sites reported not receiving feedback, which demotivated them from continuous reporting. This was consistent with the findings of a study by Hall et al in Mpumalanga, South Africa, 2009 where feedback motivated continuous reporting of ADR’s (13). If health workers lack motivation, no active detection mechanisms may be implemented to ensure identification of all DR’s and their subsequent reporting.

Timeliness of a surveillance system is a key performance measure. However, in pharmacovigilance, reporting ADR’s is preceded by immediate mitigation of the effects. Targeted spontaneous reporting (TSR) is the surveillance approach that has prescribed timelines unlike the voluntary spontaneous reporting (SR) that was being evaluated which is supposed to be part of routine practice. MCAZ acknowledged reception of ADR forms within the recommended 14-day window to facilitate causality assessments. This is contrary to findings by Bate et al in Reo de Janeiro, 2012, who identified challenges with data transmission as an impediment to the timely reception of reports in resource-constrained environments (14).

All the participants stated that it was their duty to fill the notification forms and were willing to continue participating, hence the system was acceptable. However, the majority of the health workers stated that they needed training on case detection and on how to fill the notification forms. Similar findings were reported by Pirmohamed et al in Malawi, where none of the study participants was trained on ARV pharmacovigilance and this was attributed to the high staff turnover between 2007 and 2009 (5).

Zimbabwe is compliant to the WHO minimum requirements of a functional pharmacovigilance system as stipulated by the core and complementary structural pharmacovigilance indicators (2015). The MCAZ, whose mandate is to ensure medicine safety through the institution of regulatory frameworks is compliant to WHO minimum pharmacovigilance indicators for a functional PV centre [14]. A total number of 642 reports were received by 31 December 2016, translating to 4/100 000 people, having considered a population size of 15.6 million for 2015 which is a core pharmacovigilance indicator (CP1). This is remarkable for a developing PV centre. Exercising its regulatory mandate as envisaged by PV indicator C02, the authority, out of the 10 cases of product defects that were reported, 7 products were recalled. This was commendable as if fosters compliance to set regulations.

For a reporting facility to be a functional pharmacovigilance unit, it should submit submits ≥ 10 reports annually to the pharmacovigilance centre according to WHO PV indicator P1. Only 2 centres, Wilkins and Beatrice road Hospitals fit this category in Harare city and there were a total 119 centres countrywide as at 31 December 2016 since 2012. However, in 2015 alone, only 32 health facilities submitted ADR reports countrywide. This reveals that there are facilities which were initially reported but have backtracked which is a cause for concern. There is, therefore, need to investigate what might have demotivated them from maintaining the set standards as this lowers the national performance.

Active ARV ADR detections entail enquiring from the client how they are responding to the treatment. This was found to be lacking among healthcare staff in Harare city, instead, most participants revealed that they were detecting ADR’s from client complaints. Poor ARV ADR’s detection in a city results in inaccurate quantification of the prevalence of ADR’s in the postmarket surveillance period. Reasons highlighted for poor ARV ADR detection were a lack of knowledge and training among health workers, MCAZ introduced an online reporting on 1 September 2016 but the facility remained unutilised due to lack of knowledge of its existence.

The ARV ADR surveillance system was reported to be useful although the majority lack knowledge on the surveillance system. Only one facility, Wilkins hospital had an available local database of all reported ADR’S which provided an opportunity for local utilisation of this data in programming.

The participants cited non-response by MCAZ to submitted reports as a major reason for under-reporting of ARV ADR’s. On the other hand, MCAZ indicated 100% response to all submitted reports. Further analysis revealed that although MCAZ responded to all submitted reports, the communication was conveyed through the City health directorate. Unfortunately, this communication was not being disseminated to report generating facilities. Unavailability of reporting forms, lack of appreciation of the importance of reporting ADR’s and health workers being overwhelmed with other responsibilities were other reasons attributed to under-reporting. This is contrary to the scope of accepting ADR detection as part of routine clinical practice as indicated by the same participants which also consistent with findings by Wiholm et al, 2004 in Swaziland who identified lack of appreciation of the value of ADR reporting in the post-market surveillance period as a hindrance to reporting (14).

## Conclusions

We concluded that knowledge among health workers in the city was average and the quality of data generated and committed to Vigiflow was good. City council imposed protocol of referring suspected ADR’s impeded reporting directly from facilities. Possible reasons for under-reporting ADR’s were a lack of knowledge of health workers, weak incident detection strategies, local protocol and poor information dissemination within the council. Though MCAZ were responding to the reports, their responses were not being disseminated to report generating facilities. MCAZ was fulfilling its mandate of ensuring pharmacologic safety as envisaged by the minimum PV indicator compliance as well as exercising its regulatory authority by licensing and recalling defective medicines

The ADR pharmacovigilance system was therefore found to be useful, simple, acceptable, sensitive, unstable and not representative We, therefore, recommended training of all untrained health workers involved in ADR pharmacovigilance. ADR case definitions and notification forms were distributed to health facilities in the city which did not have these. The local authority was engaged, in liaison with MCAZ, for a possible review of the local policy and facilitate reporting of ADR’s from detecting health facilities. MCAZ and Harare city directorate pledged to explore effective feedback dissemination mechanisms that will ensure all facilities receive feedback for reported ADR’s

## Acknowledgements

Special gratitude goes to my supervisors, Dr O. Mugurungi and Professor M. Tshimanga for their prodding and guidance throughout the study. I also appreciate Ms Tsitsi Juru who generously offered her assistance in the preparation of this project. I also express my gratitude to the Department of Community Medicine, University of Zimbabwe and Health Studies Office, Zimbabwe for all the help they rendered to me. Last, but not least, I would like to thank all the Masters in Public Health (MPH) colleagues for their assistance, my dear wife, son and daughter for the social support throughout the project.

